# Number of children and body composition in later life among men and women: Results from a British birth cohort study

**DOI:** 10.1101/492314

**Authors:** Charis Bridger Staatz, Rebecca Hardy

## Abstract

**Background:** Although research has found associations between increasing number of children and higher body mass index (BMI), there has been limited research investigating the association with body composition despite abdominal fat being associated with cardiovascular and metabolic risk independently of general adiposity. Most existing research has focussed on women, but investigating the relationship in men can help distinguish biological effects of pregnancy from social pathways related to parenthood.

**Methods:** Using the MRC National Survey of Health and Development (NSHD) multiple regression models were applied to test associations between number of children and body composition at age 60-64 (N=2229) and body mass index (BMI) and waist circumference (WC) at ages 60-64 and 69 (N=2149).

**Results:** In adjusted models, associations were observed between increasing numbers of children and increasing fat-adjusted lean mass index in women (p=0.06). Among men, those with children had 0.59kg (95% CI: 0.15 to 1.02) greater lean mass index than those without and fat:lean mass ratio was greater in those with 4+ children because of their slightly higher mean fat mass. Weak evidence of a higher android:gynoid mass ratio in women with children (0.03, 95% CI: 0.00,0.06, p=0.1) was observed with no associations with fat mass index or android or gynoid fat mass. Increasing BMI was observed with increasing parity in women at 60-64 and more strongly at 69 years where associations among men were also observed more clearly.

**Conclusion:** There was little evidence of a consistent association between number of children and body composition in early old age. The strongest associations are observed for lean, rather than fat mass, and in men rather than women, suggesting little evidence of biological effects of pregnancy in women. The results indicate social pathways associated with parenthood are the likely underlying mechanisms, with suggestion there may be selection into parenthood among men.

## Introduction

Characteristics of family formation such as number of children, age of first birth, age of last birth and birth intervals have been associated with subsequent overweight and obesity (1–7). The majority of studies have focussed only on women due, at least in part, to the assumption that there is a biological pathway between pregnancy, child birth and obesity. The majority of studies have found parous women to have greater body mass index (BMI), greater risk of obesity (1, 8–10) or faster weight gain trajectories (2, 3, 11, 12) than nulliparous women, with associations also observed between increasing number of children and higher BMI. However, differences in associations do exist according to the age at BMI measurement (11) with few studies investigating the relationship with BMI measured past mid-life, area of residence (non-metropolitan compared to metropolitan) (13), and according to race and ethnicity (3, 10–13). Social and behavioural factors related to parenting provide an alternative explanation to the biological effects of pregnancy for any such association. For example, parents have been found to eat more saturated fat than non-parents (12, 14), and those with 4 or more children are more likely to be smokers, inactive and from a manual social class (1). It has therefore been suggested that parenthood changes the social contexts in which people live in a way that impacts their BMI (12).

Biological pathways, such as reduced exposure to oestrogen or increased levels of relative insulin resistance (15), have been implicated in the association between pregnancy and cardiovascular disease (CVD), with the effect of pregnancy on abdominal obesity being one potential explanatory mechanism (16). Hence, biological effects of pregnancy may be particularly strong when considering fat distribution and body composition, rather than general markers of adiposity such as BMI. There is increasing evidence that fat mass and android and visceral fat are related to cardiovascular and metabolic risk factors independently of BMI and even among those who are not overweight (17–19). Few studies have assessed the association between parity and body composition. Existing studies, which have shown an association between parity and increased visceral adiposity measured using computed tomography (CT), have been based on small sample sizes (n≤170) (20, 21). One of these studies considered the change in adiposity pre- and post-pregnancy (21). The other study, based on a highly age heterogeneous sample (age 18 to 76), also considered percent body fat measured by dual-energy x-ray absorptiometry (DXA) scans finding no association with parity after adjustment for age, physical activity and smoking (20). A larger study using DXA scans to measure body composition at a mean age of 59.7 years carried out in an Asian population, did not consider number of children, but found that earlier age of first birth was associated with increased trunk fat mass (22). However, these findings may not be generalizable to western populations as visceral and abdominal fat distribution for a given BMI is higher in Asian compared to western populations (23).

There has been growing interest in the impact of becoming a father on health, with a few studies suggesting that number of children is associated with obesity and higher BMI in men as well as women (1, 9). It remains unclear as to whether associations are stronger in men or women as results have so far been inconsistent (2), and to our knowledge there have been no studies investigating associations with body composition in men. Investigating the relationship between parenthood and obesity in men as well as women can also help distinguish a biological effect of pregnancy from a social pathway related to parenthood (8); similar associations in both sexes would suggest that the underlying mechanisms are social as opposed to biological.

Using data from the Medical Research Council (MRC) National Survey of Health and Development (NSHD), a British birth cohort born in 1946, we investigated the relationship between number of children and body composition measured using DXA scans at ages 60-64 in men and women. Previous research in NSHD has shown that greater number of children in women, and being a parent in men, was associated with higher BMI at age 53. Therefore we also assess whether such associations remain into old age, investigating the association between number of children and BMI and waist circumference (WC) measured at the same age as body composition (60-64) and also at 69. We assess the extent to which associations can be explained by lifestyle and behavioural factors related to family life.

## Materials and methods

The MRC NSHD is a socially stratified sample of 2547 women and 2815 men born in one week in March 1946 in England, Scotland and Wales. There have been 24 follow ups including interviews with the mothers, assessments by school doctors and teachers during childhood, and home visits and postal questionnaires during adult life. When study members were aged 60-64, 2856 individuals were invited for assessment at one of six clinical research facilities (CRF) or to have a nurse visit them in their own home. Invitations were not sent to those who had died (n=778), who were living abroad (n=570), who had previously withdrawn from the study (n=594), or who had been lost to follow-up (n=564). Off those invited, a total of 2229 (78%) were assessed, with 1690 (59%) attending a CRF and the remaining 539 being visited at home. At the clinic visit study members underwent (DXA) scans. The most recent data collection took place in 2015 when participants were aged 69 years when 2149 were interviewed and examined in their own homes by a team of research nurses.

Ethical approval for the age 60-64 data collection was obtained from the Greater Manchester Local Research Ethics Committee and the Scotland A Research Ethics Committee, and for the age 69 data collection from the Queens Square Research Ethics Committee and the Scotland A Research Ethics Committee. Written informed consent was obtained from the study members at each data collection.

### Outcome variables

Measures of body composition were obtained at the CRFs using a QDR 4500 Discovery DXA scanner (Hologic Inc, Bedford, MA, USA) whilst the individuals were in a supine position. The measures of body composition used in these analyses were whole body, android and gynoid fat mass and whole body lean mass defined as total mass minus fat and bone mass. For all measures, mass from the head was excluded and measures were converted to kg. Full details described elsewhere (24, 25). Fat mass index (kg/m^2^) and lean mass index (kg/m^2^) were calculated by dividing the measures by height^2^. Fat-to-lean mass ratio and android-to-gynoid fat mass ratio were derived.

The anthropometric measures of height, weight, and waist circumference were measured by the nurses using standard protocols at ages 60-64 at both CRFs and home visits and at 69 at the home visits. BMI (defined as weight (kg)/height(m)^2^) was calculated.

### Explanatory variable

Number of children was derived from the number of new live births since age 53 reported at age 60-64 added to previous reports of number of live births collected throughout adult life at ages of 36, 43 and 53. For the purpose of analysis and due to the small numbers with four or more children, all those who reported having four or more children were combined into one group.

### Confounding variables

Potential confounders, identified from previous research as being associated with both number of births and adiposity, were childhood cognitive function, childhood and adult socioeconomic position, education and cigarette smoking. Cognitive function in childhood was calculated from the sum of standardised verbal and non-verbal cognitive tests taken at age 8. Socioeconomic position was measured by father’s occupational social class when the study member was aged 4 (or 7 or 11 if missing) and by occupational social class of the head of the household at age 53. The Register General classification system was used to categorise into I Professional; II Intermediate; IIINM Skilled (Non-Manual); IIIM Skilled (Manual); IV Partly skilled; V Unskilled. Highest educational qualification by age 26 was classified into (1) No Qualifications; (2) Below O-Level; (3) O-Level or Equivalent; (4) A-Level or Equivalent; (5) Degree Level or higher. Smoking status was self-reported at the age of 60-64 (and at previous ages) and was classified as current smoker, ex-smoker and never smoker.

### Statistical analysis

Multivariable regression models were used to examine the associations between number of children (with no children as the reference group) and each DXA measure and BMI and WC at 60-64. A test for deviation from a linear trend across groups of number of children was carried out and where significant deviations from linearity were detected (P≤0.05), tests for heterogeneity across groups were carried out. Where there was no evidence of deviation form linearity, number of children was fitted as a continuous variable to obtain a test for linearity. Analyses were carried out separately for men and women, and in addition, sex by number of children interactions were included to test whether the associations differed between men and women in models including both sexes. Models were then adjusted for all potential confounding variables and for lean mass index as the outcome, models were further adjusted for fat mass index due to the adaptive changes in lean mass with increasing fat mass(25). Similar models were fitted for BMI and WC at age 69 to assess whether associations were consistent with those at age 60-64. Since it has previously been observed that there are differences in outcomes between those with children and those without(1, 26), models were also fitted to test differences between those with at least one child and those with no children and then, in parents only, to test associations with number of children using one child as the reference group. All analysis was carried out using Stata14 (Stata Corp., College Station, TX, USA).

### Multiple imputation

Since there was some missing covariate information, multiple imputation was used in order to maintain the sample size. A total of 20 imputed data sets were obtained using chained equations including all variables in the adjusted models and additional variables to aid the imputation process (other measures of socioeconomic position, earlier life BMI and waist circumference). Rubin’s rule was used to combine the estimates from regression models from each of the 20 datasets. For fat mass index, lean mass index and fat-to-lean ratio, 304 (20%) (Male=152, Female=152) of the analytic sample had missing data on at least one covariate. For android fat, gynoid fat and android-gynoid ratio, the number was 318 and for BMI and waist circumference aged 60-64, 456 and 457, respectively. At age 69, the number was 348 for both BMI and waist circumference.

### Sensitivity analysis

The characteristics of those who attended a CRF, and who therefore had a DXA scan at age 60-64, have previously found to differ from those who had a home visit, with indication that they tended to be healthier(27). Therefore a sensitivity analysis was carried out to examine whether this difference in sample characteristics might have influenced associations in the subgroup who had DXA measures. We compared the difference in mean BMI and WC at age 60-64 between those who had DXA measures and those who had a home visit (and therefore did not have DXA measures). We then repeated analysis of BMI and WC at age 60-64 in the sample who had DXA measures.

## Results

Characteristics of the sample (N=2331, 52.2% female) who had at least one of the outcome variables and a valid record of number of live births are presented in Table 1. Mean fat-to-lean ratio, fat mass index and gynoid fat were higher in women compared to men, whilst all other measures were higher in men. Mean BMI was similar in men and women at both ages 60-64 and 69 and, as expected, mean WC was higher in men compared to women. The median number of children was 2 for both men and women, with few individuals with 4 or more children (7.27% and 8.46%, respectively).

**Table 1.** Characteristics of men and women in the analytic sample.

| Variable | Male |  | Female |  |
| --- | --- | --- | --- | --- |
|  | *Number (%)* | *Mean (SD)* | *Number (%)* | *Mean (SD)* |
| Fat Mass Index at 60-64 (kg/m<sup>2</sup>) | 701 | 7.72 (2.33) | 787 | 11.1 (3.47) |
| Lean Mass Index at 60-64 (kg/m<sup>2</sup>) | 701 | 17.42 (1.97) | 787 | 14.18 (1.84) |
| Fat- Lean Mass ratio at 60-64 | 701 | 0.44 (0.11) | 787 | 0.77 (0.19) |
| Android Fat at 60-64 (kg) | 731 | 2.49 (0.97) | 830 | 2.33 (1.02) |
| Gynoid Fat at 60-64 (kg) | 731 | 3.76 (1.01) | 830 | 5.13 (1.48) |
| Android-Gynoid Ratio at 60-64 | 731 | 1.18 (0.18) | 830 | 0.92 (0.15) |
| Body Mass Index at 60-64 years (kg/m<sup>2</sup>) | 990 | 27.85 (4.08) | 1,117 | 28.00 (5.51) |
| Waist Circumference at 60-64 years (cm) | 990 | 100.68<br>(11.02) | 1,115 | 92.56 (13.09) |
| Body Mass Index at 69 years (kg/m<sup>2</sup>) | 961 | 28.15 (4.55) | 1,047 | 28.21 (5.96) |
| Waist Circumference at 69 years (cm) | 961 | 100.93<br>(12.21) | 1,053 | 92.39 (13.91) |
| **Number of Children\*** |  |  |  |  |
| 0 | 163 (14.63) |  | 133 (10.93) |  |
| 1 | 117 (10.50) |  | 142 (11.67) |  |
| 2 | 551 (49.46) |  | 563 (46.26) |  |
| 3 | 202 (18.13) |  | 276 (22.68) |  |
| 4+ | 81 (7.27) |  | 103 (8.46) |  |
| **Total** | 1,114 |  | 1,217 |  |

\*Taken from the number of children reported at age 60-64

### Body composition

Fat mass index and lean mass index were available for 701 men and 787 women and android and gynoid fat for 731 men and 830 women. There was little evidence of a trend across number of children categories for the majority of body composition measures in either men or women (Table 2). Associations between number of children and lean mass index were observed for both sexes. Higher mean lean mass index was observed in men with children compared to those without and there was weak evidence that increasing parity was associated with increasing lean mass index in women (although there was no evidence of a sex interaction). These findings were confirmed in the analysis comparing men with one or more children to those with no children where the mean difference in lean mass index was 0.59kg (95% CI: 0.15 to 1.02, P=0.008) and among those with at least one child for women (Table.3). Among men, there was also a suggestion of a non-linear relationship between number of children and fat:lean mass ratio where those with 1, 2 or 3 children had lower ratios than men with either 0 or 4+ children (Table 2). Men with 4+ children, despite having greatest lean mass index, also had greatest mean fat mass index, resulting in a ratio which was greater than men with fewer children. The higher fat mass in men with 4+ children was confirmed in analyses including only parents (p=0.05, Table.3). Associations were hardly affected following adjustment for confounders (Table.2) with slight attenuation of effects for lean mass index and very slight strengthening for the ration in men. Among women, there was weak evidence that those with children had higher android:gynoid fat mass ratio than those without children (0.03 (95% CI: 0.00-0.06) higher, p=0.1) (Table.3).

**Table 2.** Adjusted and unadjusted regression coefficients (mean differences) and 95% confidence intervals (95% CI) for each measure of body composition at 60-64 years and body mass index and waist circumference at 60-64 and 69 years in men and women.

| Outcome Variable | N | Unadjusted |  |  |  |  | Adjusted** |  |  |  |  |
| --- | --- | --- | --- | --- | --- | --- | --- | --- | --- | --- | --- |
|  |  | Regression coefficients (95% CI) compared with baseline group of 0 children |  |  |  |  | Regression coefficients (95% CI) compared with baseline group of 0 children |  |  |  |  |
|  |  | 1 | 2 | 3 | 4+ | P value, trend | 1 | 2 | 3 | 4+ | P value, trend |
| Fat Mass Index (kg/m2) |  |  |  |  |  |  |  |  |  |  |  |
| Male | 701 | 0.17<br>(-0.55,0.91) | -0.02<br>(-0.55,0.50) | -0.33<br>(-0.95,0.29) | 0.69<br>(-0.11,1.48) | 0.8 | -0.05<br>(-0.79,0.69) | -0.02<br>(-0.56,0.52) | -0.43<br>(-1.05,0.20) | 0.64<br>(-0.17,1.45) | 0.9 |
| Female | 787 | 0.66<br>(-0.33, 1.64) | 0.13<br>(-0.65,0.91) | 0.42<br>(-0.45,1.29) | 0.75<br>(-0.33,1.83) | 0.3 | 0.38<br>(-0.62,1.38) | -0.03<br>(-0.82,0.76) | 0.23<br>(-0.65,1.10) | 0.42<br>(-0.68,1.53) | 0.6 |
| Lean Mass Index (kg/m2) |  |  |  |  |  |  |  |  |  |  |  |
| Male | 701 | 0.98<br>(0.37,1.60) | 0.56<br>(0.11,1.00) | 0.56<br>(0.03,1.08) | 1.02<br>(0.35,1.69) | 0.008* | 0.85<br>(0.23,1.48) | 0.53<br>(0.08,0.98) | 0.47<br>(-0.06,0.99) | 1.05<br>(0.36,1.74) | 0.02* |
| Female | 787 | 0.14<br>(-0.39,0.66) | 0.03<br>(-0.39,0.44) | 0.34<br>(-0.12,0.80) | 0.42<br>(-0.15,0.99) | 0.07 | 0.10<br>(-0.43,0.64) | 0.03<br>(-0.39,0.46) | 0.33<br>(-0.13,0.80) | 0.39<br>(-0.20,0.98) | 0.08 |
| Fat: Lean Mass ratio |  |  |  |  |  |  |  |  |  |  |  |
| Male | 701 | -0.01<br>(-0.05,0.02) | -0.02<br>(-0.04,0.01) | -0.03<br>(-0.06,0.00) | 0.01<br>(-0.03,0.05) | 0.07* | -0.02<br>(-0.06,0.01) | -0.02<br>(-0.04,0.01) | -0.04<br>(-0.07,-0.01) | 0.01<br>(-0.03,0.05) | 0.05* |
| Female | 787 | 0.05<br>(0.00,0.10) | 0.02<br>(-0.03,0.06) | 0.02<br>(-0.03,0.06) | 0.04<br>(-0.02,0.09) | 0.7 | 0.03<br>(-0.02,0.08) | 0.00<br>(-0.04,0.05) | 0.00<br>(-0.04,0.05) | 0.01<br>(-0.05,0.7) | 0.9 |
| Android Fat (kg) |  |  |  |  |  |  |  |  |  |  |  |
| Male | 731 | 0.09<br>(-0.21,0.39) | 0.02<br>(-0.19,0.24) | -0.08<br>(-0.33,0.17) | 0.24<br>(-0.08,0.56) | 0.7 | -0.01<br>(-0.31,0.29) | 0.00<br>(-0.21,0.22) | -0.14<br>(-0.39,0.11) | 0.21<br>(-0.11,0.54) | 0.9 |
| Female | 830 | 0.16<br>(-0.12,0.45) | 0.07<br>(-0.15,0.30) | 0.15<br>(-0.10,0.39) | 0.25<br>(-0.06,0.56) | 0.2 | 0.12<br>(-0.17,0.41) | 0.05<br>(-0.17,0.28) | 0.12<br>(-0.13,0.38) | 0.22<br>(-0.10,0.54) | 0.2 |
| **Gynoid Fat (kg)** |  |  |  |  |  |  |  |  |  |  |  |
| *Male* | 731 | 0.14<br>(-0.17,0.45) | 0.02<br>(-0.21,0.24) | -0.01<br>(-0.27,0.26) | 0.31<br>(-0.02,0.65) | 0.4 | 0.08<br>(-0.24,0.40) | 0.01<br>(-0.22,0.24) | -0.03<br>(-0.30,0.24) | 0.31<br>(-0.04,0.66) | 0.4 |
| *Female* | 830 | 0.10<br>(-0.31,0.52) | -0.09<br>(-0.42,0.24) | 0.00<br>(-0.36,0.36) | 0.20<br>(-0.25,0.65) | 0.7 | 0.04<br>(-0.39,0.46) | -0.11<br>(-0.44,0.22) | 0.02<br>(-0.39,0.34) | 0.15<br>(-0.31,0.62) | 0.8 |
| **Android/Gynoid Ratio** |  |  |  |  |  |  |  |  |  |  |  |
| *Male* | 731 | -0.00<br>(-0.06,0.05) | 0.01<br>(-0.03,0.05) | 0.00<br>(-0.05,0.05) | 0.01<br>(-0.05,0.7) | 0.9 | -0.01<br>(-0.07,0.05) | 0.01<br>(-0.03,0.05) | -0.01<br>(-0.06,0.04) | 0.01<br>(-0.05,0.08) | 0.9 |
| *Female* | 830 | 0.03<br>(-0.01,0.07) | 0.03<br>(0.00,0.07) | 0.02<br>(-0.01,0.06) | 0.03<br>(-0.01,0.08) | 0.2 | 0.02<br>(-0.02,0.07) | 0.03<br>(0.00,0.06) | 0.02<br>(-0.01,0.06) | 0.03<br>(-0.01,0.08) | 0.2 |
| **Body Mass Index (kg/m2)** |  |  |  |  |  |  |  |  |  |  |  |
| **60-64** |  |  |  |  |  |  |  |  |  |  |  |
| *Male* | 990 | 0.77<br>(-0.26,1.80) | 0.48<br>(-0.27,1.22) | 0.36<br>(-0.52,1.25) | 1.34<br>(0.17,2.51) | 0.1 | 0.43<br>(-0.60,1.45) | 0.38<br>(-0.36,1.13) | 0.15<br>(-0.72,1.03) | 1.26<br>(0.09,2.42) | 0.3 |
| *Female* | 1,117 | 0.44<br>(-0.91,1.80) | 0.35<br>(-0.74,1.43) | 1.19<br>(0.00,2.38) | 1.43<br>(-0.04,2.91) | 0.02 | 0.13<br>(-1.23,1.50) | 0.11<br>(-0.98,1.21) | 0.87<br>(-0.33,2.07) | 1.07<br>(-0.42,2.56) | 0.05 |
| **Waist Circumference (cm)60-64** |  |  |  |  |  |  |  |  |  |  |  |
| *Male* | 990 | 0.99<br>(-1.80,3.78) | 0.39<br>(-1.63,2.42) | 0.49<br>(-1.91,2.88) | 2.87<br>(-0.28,6.00) | 0.3 | -0.14<br>(-2.91,2.62) | 0.09<br>(-1.92,2.12) | -0.03<br>(-2.41,2.35) | 2.41<br>(-0.73,5.55) | 0.3 |
| *Female* | 1,115 | 1.56<br>(-1.66,4.79) | 0.61<br>(-1.97,3.19) | 1.30<br>(-1.53,4.13) | 4.34<br>(0.82,7.86) | 0.06 | 0.77<br>(-2.47,4.02) | -0.04<br>(-2.64,2.56) | 0.48<br>(-2.37,3.33) | 3.54<br>(-0.02,7.11) | 0.2 |
| **Body Mass Index (kg/m2)** |  |  |  |  |  |  |  |  |  |  |  |
| **69** |  |  |  |  |  |  |  |  |  |  |  |
| *Male* | 961 | 1.46<br>(0.26,2.65) | 0.37<br>(-0.53,1.27) | 0.75<br>(-0.29,1.80) | 1.92<br>(0.58,3.26) | 0.01* | 1.07<br>(-0.11,2.26) | 0.31<br>(-0.58,1.21) | 0.58<br>(-0.46,1.62) | 1.69<br>(0.34,3.03) | 0.07* |
| *Female* | 1,047 | 0.23<br>(-1.22,1.68) | 0.24<br>(-0.92,1.40) | 1.06<br>(-0.20,2.33) | 2.19<br>(0.61,3.78) | 0.003 | -0.11<br>(-1.56,1.34) | -0.13<br>(-1.30,1.04) | 0.71<br>(0.-0.57,1.98) | 2.68<br>(0.08,3.29) | 0.02 |
| **Waist Circumference** |  |  |  |  |  |  |  |  |  |  |  |
| **(cm) 69** |  |  |  |  |  |  |  |  |  |  |  |
| *Male* | 961 | 2.55<br>(-0.66,5.77) | 0.57<br>(-1.85,2.99) | 1.34<br>(-1.46,4.14) | 5.97<br>(2.37,9.57) | 0.006* | 1.35<br>(-1.86,4.55) | 0.25<br>(-2.17,2.67) | 0.85<br>(-1.95,3.65) | 5.09<br>(1.47,8.70) | 0.04* |
| *Female* | 1,053 | -0.17<br>(-3.72,3.36) | 0.13<br>(-2.70,2.97) | 2.00<br>(-1.10,5.10) | 3.70<br>(-0.17,7.58) | 0.02 | -1.33<br>(-4.88,2.21) | -1.00<br>(-3.86,1.86) | 0.87<br>(-2.24,3.98) | 2.25<br>(-1.67,6.17) | 0.1 |

\*Test for heterogeneity across groups when test for deviation from linearity gives $P \leq 0.05$ .

**Table.3.**
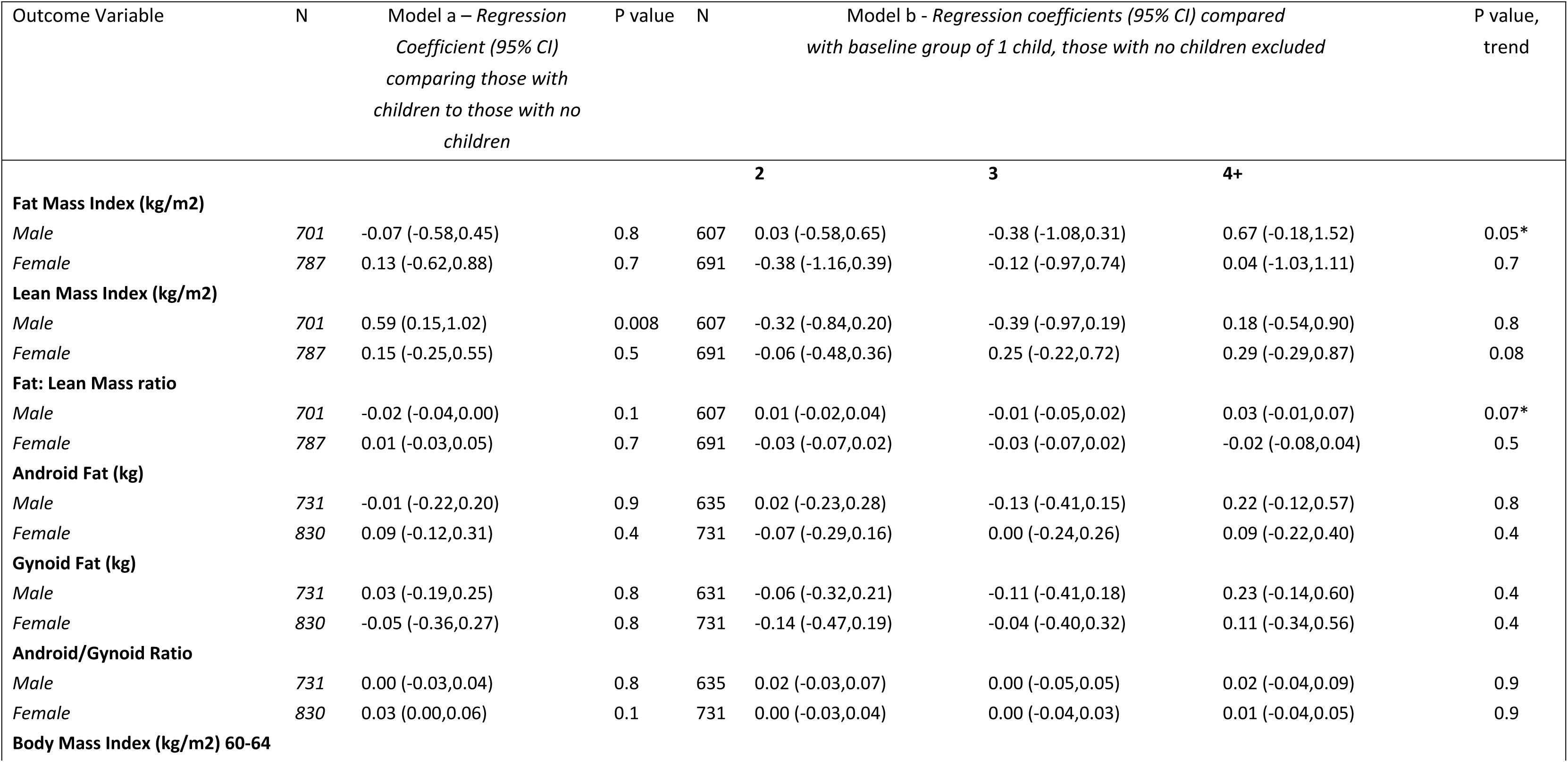
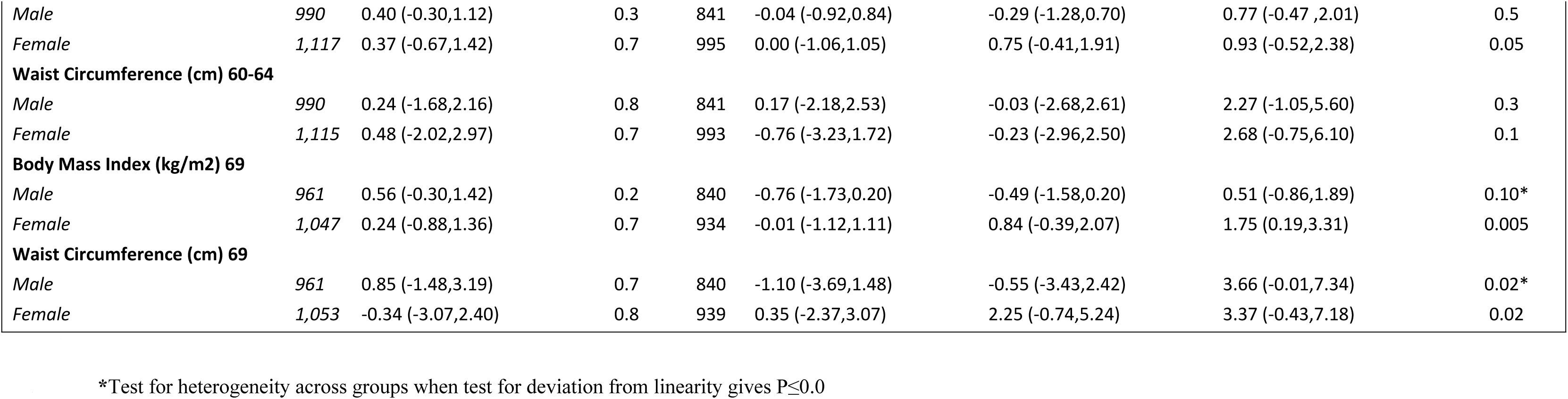
Adjusted (adjusted model includes childhood cognitive function, childhood and adult socioeconomic position, education and cigarette smoking) regression coefficients (mean differences) and 95% confidence intervals (95% CI) for each measure of body composition and anthropometric measures aged 60-64 and 69 for (a) parents versus non-parents, and (b) number of children for parents only – reference group 1 child.

The associations with lean mass index for men and women remained after adjustment for fat mass index. (Table.4).

**Table 4.**
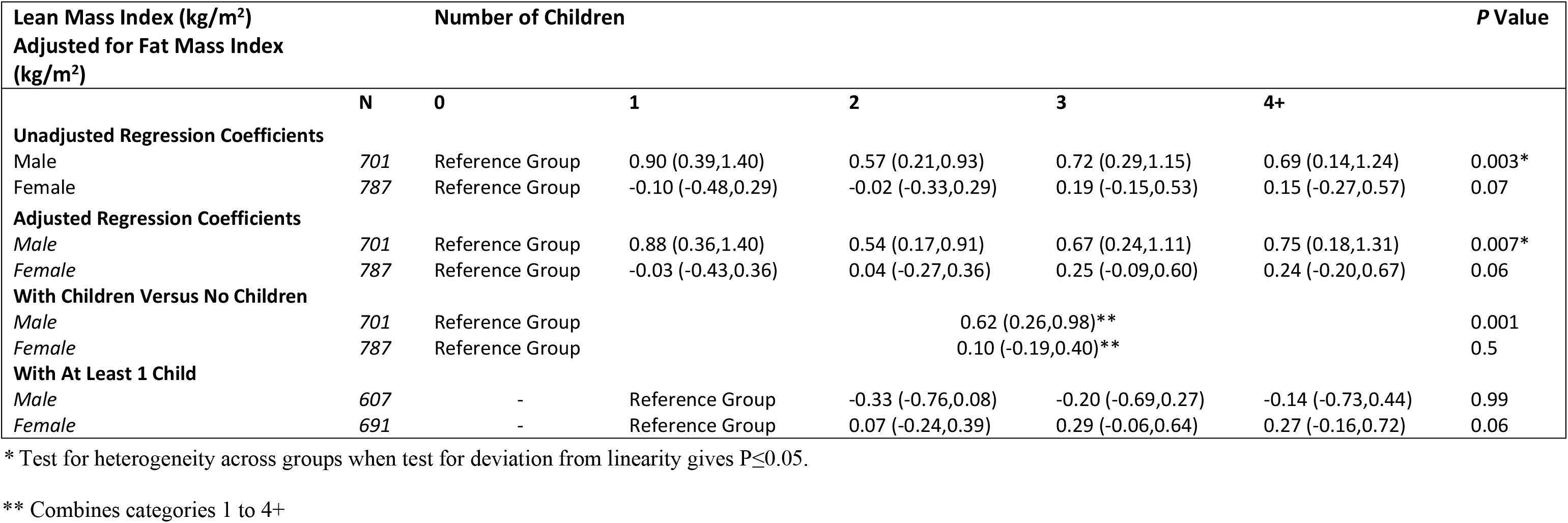
Regression models for lean mass index (kg/m^2^) adjusted for fat mass index (kg/m^2^).

### BMI and WC

BMI at age 60-64 increased with increasing parity, with the highest regression coefficients among women with 3 or 4+ children. For WC the trend was weaker, although those with 4+ children had the highest mean (Table.2). There is no evidence of a difference between those with children and those without for either BMI and WC with weak increasing trends among parous women. Among men, no evidence of trends across groups were observed, although those with 4+ children had higher mean BMI and WC than other groups. No significant sex interactions were observed for either BMI or WC (Table.2). Linear associations for both outcomes are attenuated in women following adjustment, but remain significant for BMI (p=0.05, Table.2).

The trends of increasing BMI and WC with increasing number of children in women are stronger at age 69 than at age 60-64. In men, associations are also observed at 69 years with mean levels highest in those with 4+ children (Table.2). Associations are slightly attenuated in adjusted models.

### Sensitivity analysis

The mean BMI and WC at 60-64 were lower in men and women who had a DXA scan compared with those who did not (two tailed t-test, P<0.001). Regression analysis showed weaker associations between number of children and both BMI and WC in the unadjusted and adjusted model for those that attended the DXA compared to the whole sample (S1. Table).

## Discussion

There was no clear evidence of a consistent association between number of children and body composition in early old age in either men or women and no evidence of clear sex differences in associations. Lean mass index was higher in men with children than those without children, while men with 4+ children had higher fat-lean ratio than others. In women, there was weak evidence of increasing lean mass index with increasing number of children and higher android:gynoid fat mass ratio in those with compared to without children. Both BMI and WC at ages 60-64 and, more clearly, at 69 years increased with increasing numbers of children in women, whilst highest mean BMI and WC was observed among men with 4+ children, particularly at 69 years.

Limited previous research has investigated the association between number of children and body composition, particularly in men or at older ages. The weak association between having children and higher android:gynoid fat mass ratio could be consistent with existing research showing increases in visceral fat measured using computed tomography (CT) scans, although not fat mass measured by DXA, among 14 women who had a child within a 5 year interval compared with 108 who did not(21). Another small study (n=170) in an age-heterogeneous group of women from 18-70 years found that after adjustment, increasing number of children was associated with increasing intra-abdominal adipose tissue but not with total fat mass(20). Other research in a larger sample of women from Korea with body composition measured at older ages (45 years and older) has shown younger age at first birth (those with younger age at first birth tend to have a higher number of children) to be associated with higher total and truncal fat mass measured using DXA(22).

Our finding of associations, particularly among women, between number of children and BMI are more consistent with previous findings. In previous research carried out using the NSHD cohort at age 53, BMI from age 36 was found to increase with increasing number of children in both men and women with stronger associations in women, although associations in women showed greater attenuation once adjusted for social factors, and no sex interactions were identified (1). A study by Lawlor et al (2002) also found linear associations in men and women for both BMI and WHR, with stronger associations observed in women and attenuation to the null for WHR in men once adjusted for lifestyle risk factors and SEP, but no attenuation in women. Two US studies found no evidence that number of children increased weight gain more in women than in men (9, 10) with one even finding greater weight gain among men when stratified according to SEP and race (10).

To our knowledge no other studies have considered associations with lean mass, the outcome where we observed the strongest associations. That we found higher mean lean mass index in men who had children compared to those who did not and, more weakly, increasing lean mass index with increasing parity in women could indicate a degree of selection into parenthood. This effect could be greater for men, this is consistent with previous research in NSHD finding that physical performance, including grip strength which is an indicator of muscle strength, was lower among unmarried and childless men compared to others (26). An alternative explanation for the greater muscle mass among those with children is that with increasing fat mass, muscle mass also increases to accommodate carrying the extra weight (25), although the associations remained once models were adjusted for fat mass index. It is possible that this selection could simultaneously occur with a second process of accumulation of fat associated with high numbers of children given the higher fat: lean mass ratio among men with 4+ children.

Our findings suggest that for women the effects of pregnancy on body composition in older age are likely to be small. While the similar associations in men and women for BMI and WC, which are somewhat attenuated after adjustment instead, suggest that the changing social contexts associated with parenthood for both sexes are likely to be key in explaining those associations. Our research extends previous findings by showing that such associations extend into old age. It has been found that the diets of parents contain more saturated fat than non-parents (14, 28, 29) and an increased intake of sugar sweetened beverages and total energy (28). Physical activity has also been identified as being lower in parents compared to non-parents (28–32). Some research has found that inactivity is associated with increased parity in women (31) whilst others have found that motherhood itself, as opposed to number of children, is key in explaining reduced levels of physical activity (30).

Barriers to maintaining physical activity such as guilt, family responsibilities, lack of support, scheduling constraints, and work have been identified as reasons for the difference in activity patterns between parents and non-parents (33). Some studies have even found that identities formed around being a parent contradict positive health behaviours (34) as there is a shift from prioritising individual needs to that of their children and family (33, 34). Where parents have been able to maintain physical activity after childbirth, exercise levels before and during pregnancy, exercise self-efficacy (35, 36) and the ability to develop strategies to incorporate exercise into family life (33) have been highlighted. Additional possible behaviour changes in parents relate to smoking and drinking, with research focusing solely on mothers finding both increased levels of smoking (29, 31) and drinking (31) compared to non-mothers.

There are some limitations with the current study. Firstly, body composition was measured only at age 60-64 meaning that associations with the number of children could not be assessed closer to age of childbearing in order to see whether the associations change with increasing age. Moreover, the use of DXA scans do not provide detailed insight to levels of visceral compared to subcutaneous fat which may be more relevant for pregnancy. However, research has shown that DXA is a suitable method for assessing body composition, with high correlations between android fat measured by DXA scans and visceral fat measured by CT scans (37, 38). Additionally if the effects of number of children on outcomes are small, and greatest at high parity, our study may lack power to investigate the effect of high numbers of children due to few individuals with 4 or more children. However, the study had the power to observe associations with BMI.

There is possibility of selection bias through loss to follow up if those individuals who have dropped out of the study or died differ in their relationship between number of children and the outcome variables. Further, the sample who attended clinics for DXA assessment have previously been shown to be healthier than those who participated in home visits (and who had basic anthropometric measures but not DXA). The sensitivity analysis provided some evidence to suggest that associations may be weaker in the sample who attended clinics as the association with BMI was weaker in this group.

Our findings are from a sample born in Britain 1946 (39) and although the cohort is nationally representative of the population at the time of recruitment, it does not include immigrants into Britain of the same age and may not be representative of later generations. Research suggests that more recent cohorts are experiencing the obesogenic environment earlier in the life course (40) and childbearing patterns have changed such that on the whole couples have later and fewer births, therefore associations may vary between cohorts. Nevertheless, adjustment of social factors relevant to the cohort under study would be expected to result in a consistent biological effect of pregnancy, if one exists.

The current research is, to the best of our knowledge, unique in that it uses DXA imaging to measure body composition, rather than relying on more general anthropometric measures which cannot distinguish fat from lean mass, in a large western sample and which includes men as well as women. The current study also measures body composition as well as BMI and WC into older age.

Our findings suggest little evidence of a biological effect of pregnancy in women on body composition in older age. Social pathways, contexts and behaviours associated with parenthood are more likely to explain the persisting associations observed with more general measures of adiposity, BMI and WC, and selection into parenthood may explain associations with lean mass, particularly in men. As parents have been recognised as important in the development of health behaviours, such as physical activity engagement (41), in their children, these pathways could also have knock on effect on children’s propensity to overweight. Therefore, public health initiatives should focus on adapting the social contexts of parenthood and the health behaviours in families, as opposed to focusing on the timing and processes of pregnancy.

## Acknowledgements

We thank NSHD study members for their lifelong participation and past and present members of the NSHD study team who helped to collect the data. Data used in this publication are available to bona fide researchers upon request to the NSHD Data Sharing Committee via a standard application procedure. Further details can be found at http://www.nshd.mrc.ac.uk/data. doi:10.5522/NSHD/Q101; doi:10.5522/NSHD/Q102 CBS is supported by a PhD studentship from the UK Medical research Council (MR/N013867/1). The MRC NSHD and RH are supported by the UK Medical Research Council (MC_UU_12019/1, MC_UU_12019/2).

## Supporting information captions

**S.1 Table. Sensitivity Analysis - unadjusted and adjusted regressions for anthropometric measures aged 60-64 comparing full sample (those who had a home visit as well as those who attended the CRF) with only those who attended the CRF and thus also had a DXA scan**. ** Adjusted model includes childhood cognitive function, childhood and adult socioeconomic position, education and cigarette smoking.

